# Transfer Learning for Cognitive Reserve Quantification

**DOI:** 10.1101/2022.02.03.479059

**Authors:** Xi Zhu, Yi Liu, Christian G. Habeck, Yaakov Stern, Seonjoo Lee, the Alzheimer’s Disease Neuroimaging Initiative

## Abstract

Cognitive reserve (CR) has been introduced to explain individual differences in susceptibility to cognitive or functional impairment in the presence of age or pathology. We developed a deep learning model to quantify the CR as residual variance in memory performance using the structural MRI data from a lifespan healthy cohort. The generalizability of the sMRI-based deep learning model was tested in two independent healthy and Alzheimer’s cohorts using transfer learning framework.

Structural MRIs were collected from three cohorts: 495 healthy adults (initially aged 20-80) from RANN, 620 healthy participants (age 36-100) from lifespan Human Connectome Project Aging (HCPA), and 941 subjects from Alzheimer’s Disease Neuroimaging Initiative (ADNI). Region of interest (ROI)-specific cortical thickness and volume measures were extracted using the Desikan-Killiany Atlas. Cognitive reserve was quantified by residuals which subtract the predicted memory from the true memory. Cascade neural network (CNN) models were used to train RANN dataset for memory prediction. Transfer learning was applied to transfer the T1 imaging-based model from source domain (using RANN) to the target domain (HCPA or ADNI).

The CNN model trained on the RANN dataset exhibited strong linear correlation between true and predicted memory based on the chosen T1 cortical thickness and volume predictors. In addition, the model generated from healthy lifespan data (RANN) was able to generalize to an independent healthy lifespan data (HCPA) and older demented participants (ADNI) across different scanner types. The estimated CR was correlated with CR proxies such education and IQ across all three datasets.

The current findings suggest that the transfer learning approach is an effective way to generalize the residual-based CR estimation. It is applicable to various diseases and may flexibly incorporate different imaging modalities such as fMRI and PET, making it a promising tool for scientific and clinical purposes.

**Highlight:**

- Quantification of cognitive reserve using brain measures for pre-symptomatic Alzheimer’s patients can be estimated by leveraging lifespan data.
- Multi-center, multi-scanner, multi-sequence can affect the performance of the quantification.
- Leveraging lifespan data from a single site can improve the performance.
- Transfer learning allows the pre-trained network to successfully reconstruct the dataset acquired from different domains or age groups.

## Introduction

Approximately 15–20% of adults aged 65 or older suffer from significant cognitive decline resulting in mild cognitive impairment (MCI); among these, 11.3% adults later develop dementia due to Alzheimer’s disease (AD) (Association, 2021). With the lack of an effective treatment strategy, there is a great need to identify factors that can slow the progression to dementia and maintain quality of life (Zissimopoulos, Crimmins, & St Clair, 2014).

Cognitive reserve (CR) has been introduced to explain individual differences in susceptibility to cognitive or functional impairment in the presence of age or disease-related brain changes (Stern, 2002). Individuals with high CR have greater resilience and maintained normal cognitive function longer when confronted with late-life neuropathology. Typical CR proxy measures include years of education (Meng & D’Arcy, 2012; Stern et al., 1994), premorbid IQ (Alexander et al., 1997), occupational achievement, and engagement in cognitively and socially stimulating activities (Scarmeas & Stern, 2003). These are thought to protect against functional impairment by promoting the ability to better compensate for brain changes. However, these proxy variables of cognitive reserve is with controversy. Specifically, these proxy measures fail to provide the entirety of the construct, the same value of a proxy variable may reflect different experiences across people. In addition, most proxy measures only represent static cognitive reserve, and cannot be changed over time for adjusting the change of cognitive reserve. Lastly, these measures rely on recollection of prior activities which are an indirect proxy of CR. (Borenstein, Copenhaver, & Mortimer, 2006; Jones et al., 2011; Satz, Cole, Hardy, & Rassovsky, 2011). To address the above mentioned limitation, a direct measure of CR based on unbiased information is highly needed. One popular approach to quantify CR is to measure the residual variance between predicted cognitive performance based on individual’s level of brain status and neuropathology and the actual individual’s performance (Reed et al., 2010). This residual-based measures offer a more precise measurement of CR (Bocancea et al., 2021). High-reserve individuals exhibit higher actual measured cognitive performance than that predicted.

Structural magnetic resonance imaging (sMRI) has been used as a measure of the regional brain atrophy underlying cognitive decline and dementia (Mueller, Schuff, & Weiner, 2006). Previous studies using the sMRI to calculate the residual variance operational measure of CR showed promising results (Reed et al., 2010; Zahodne et al., 2015; Zahodne et al., 2013) in older participants. Currently, most research in cognitive aging has used life-span data (Razlighi, Habeck, Barulli, & Stern, 2017; Salthouse, 2010; Taylor et al., 2017; Tucker-Drob, 2019). However, leveraging life-span brain and cognition data in quantifying cognitive reserve has not been done.

In addition, brain imaging data from multi-sites may have high variability due to different MRI sequences of different scanners, thus, limiting direct application of a previously trained model to new datasets acquired from different sites.

Traditional machine learning methods to mitigate the influence of variability across sites require a balanced sample from each site and assume the same distribution across training and test datasets. The performance of a predictive model declines when these assumptions are violated. Transfer learning is a machine learning technique that utilizes the knowledge gained from one task and applies it to a different but related task. It is a popular optimization approach that allows rapid progress or improved performance when modeling the second task. The sMRI obtained from various sites or scanners may represent similar brain properties but may exhibit different observational distributions. Thus, the transfer learning approach may be applied to improve the generalizability of the sMRI-based residual models.

In this study, we proposed a CR quantification framework that leverages a single-site, large scale lifespan data and uses transfer learning to handle scanner and site differences. We first built a deep learning model to quantify the CR as residual variance in memory performance using the sMRI data from a healthy lifespan cohort (age 20-80). Our first goal was to assess whether using lifespan data of heathy individuals, which shows more variability in cognition function, may enable better quantification of the relationship between sMRI and cognitive performance. Second, to test the generalizability of the sMRI-based deep learning model, we utilized the transfer learning approach to fine-tune the pre-trained deep learning model to an independent, healthy lifespan cohort: the Human Connectome Project-Aging cohort (HCPA). Third, to test whether the model generated from healthy lifespan data could generalize to older MCI or demented individuals, we used transfer learning again to fine-tune the model to fit data from participants in the Alzheimer’s Disease Neuroimaging Initiative (ADNI). The ADNI datasets were acquired from different scanners and under different imaging conditions, so we could test whether the model is affected by different scanners. To validate our operationalization of CR in all three cohorts, we hypothesized that the estimated CR would correlate with education and IQ (i.e., a well-established CR proxy).

## Method and Material

### Participants

#### CR/RANN

495 healthy adults (initially aged 20-80) were drawn from our ongoing studies at Columbia University Irving Medical Center: The Reference Ability Neural Network (RANN) study and the Cognitive Reserve (CR) study (Stern, 2009; Stern et al., 2014). Demographic characteristics of these participants are summarized in Table 1.

**Table 1.** Demographic Characteristics in RANN, HCPA, and ADNI Study

|  | **RANN** | **HCPA** | **ADNI** | **p** |
| --- | --- | --- | --- | --- |
| **Total N** | 495 | 620 | 941 |  |
| **Age** |  |  |  | <0.001 |
| - Mean(SD) | 53.42 (16.90) | 59.499 (15.171) | 72.40 (7.21) |  |
| - Median(Q1,Q3) | 60.00 (38.00,67.00) | 58.04 (46.92,70.98) | 72.00 (67.00,77.00) |  |
| **Sex, n(%)** |  |  |  | 0.0065 |
| - Female | 282 (57%) | 358 (57.6%) | 474 (50.4%) |  |
| - Male | 213 (43.0%) | 263 (42.4%) | 467 (49.6%) |  |
| **Memory** |  |  |  | <0.001 |
| - Mean(SD) | 0.03(0.94) | 0.01 (0.72) | 0.45 (0.91) |  |
| - Median(Q1,Q3) | 0.13(-0.62,0.80) | -0.34 (-0.38, 0.37) | 0.53 (-0.19,1.12) |  |
| **Education** |  |  |  | <0.001 |
| - Mean(SD) | 16.20 (2.35) | 17.46 (2.19) | 16.42 (2.53) |  |
| - Median(Q1,Q3) | 16.00 (14.00,18.00) | 18.00 (16, 19) | 16 (15.00,18.00) |  |
| **IQ** | **NART IQ** | **Nih fluidcogcomp** | **NART IQ** | 0.512 |
| - N-Miss | 5 | 2 | 17 |  |
| - Mean(SD) | 117.02 (8.68) | 120.85 (139.17) | 116.35 (11.23) |  |
| - Median(Q1,Q3) | 119.20<br>(111.92, 124.00) | 100<br>(91.00, 108.00) | 119.44<br>(110.76,124.40) |  |
| **People** |  |  |  |  |
| - N-Miss | 113 | - | - |  |
| - Mean(SD) | 4.77 (2.28) | - | - |  |
| - Median(Q1,Q3) | 6.00 (3.00, 6.00) | - | - |  |
| **Data** |  |  |  |  |
| - N-Miss | 113 | - | - |  |
| - Mean(SD) | 1.73 (1.44) | - | - |  |
| - Median(Q1,Q3) | 1.00 (1.00, 3.00) | - | - |  |
| **Things** |  |  |  |  |
| - N-Miss | 113 | - | - |  |
| - Mean(SD) | 5.40 (2.40) | - | - |  |
| - Median(Q1,Q3) | 7.00 (2.00, 7.00) | - | - |  |
| **Diagnosis, n(%)** |  |  |  |  |
| CN | 495 (100%) | 620 (100%) | 417 (44.3%) |  |
| MCI | - | - | 378 (40.2%) |  |
| AD | - | - | 146 (15.5%) |  |

**Table 2.**
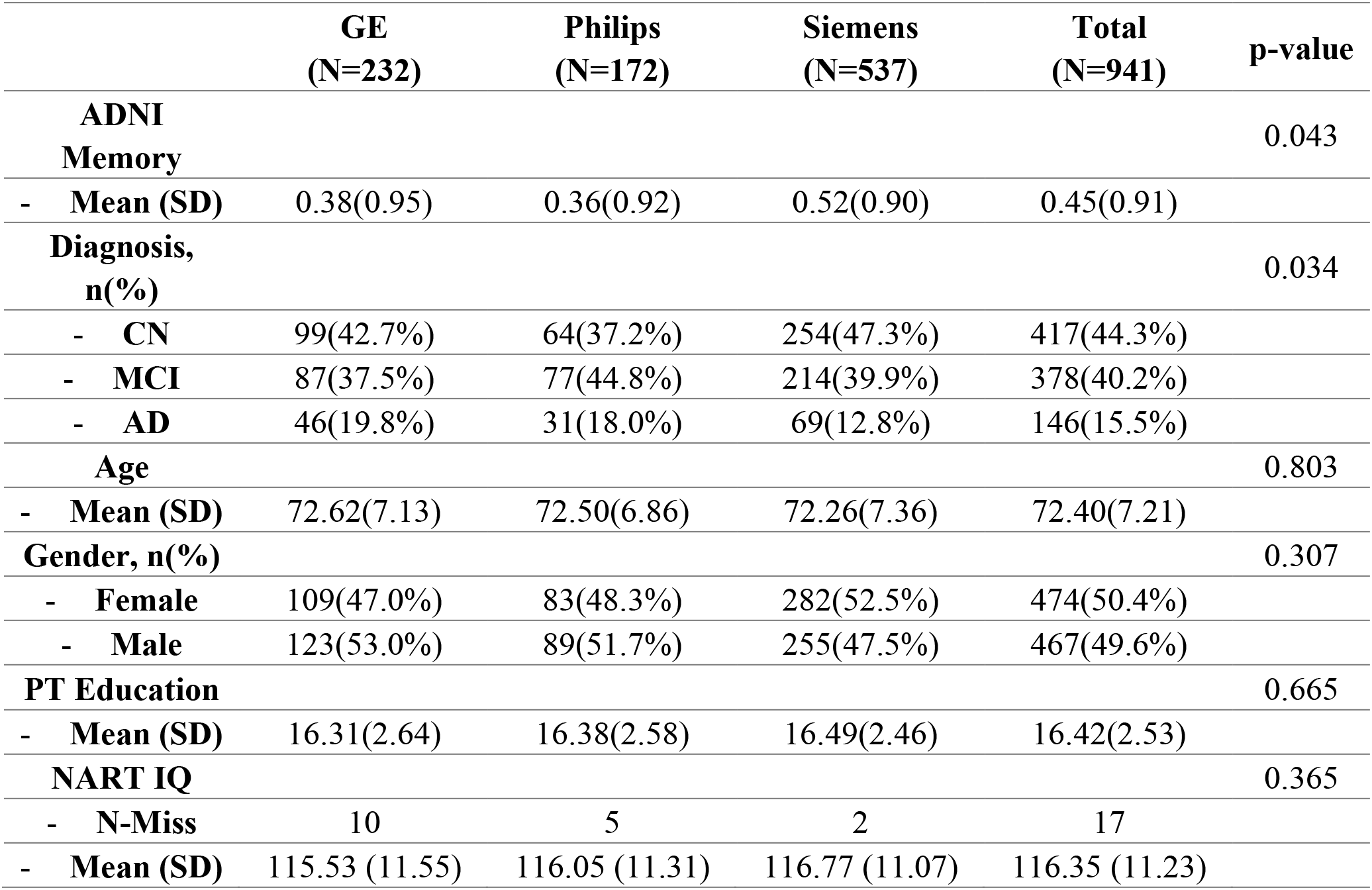
Demographic Characteristics in ADNI dataset across three scanners

Subjects were recruited primarily by randomized market mailing. An initial telephone screening determined whether participants met basic inclusion criteria (i.e., right-handed, English speaking, no psychiatric or neurological disorders, and normal or corrected-to-normal vision). Potentially eligible participants were further screened in person with structured medical and neuropsychological evaluations to ensure that they had no neurological or psychiatric conditions, cognitive impairment, or contraindication for MRI scanning. Global cognitive functioning was assessed with the Mattis Dementia Rating Scale (Lucas et al., 1998), on which a minimum score of 130 was required for retention in the study. In addition, participants who met diagnostic criteria for MCI were excluded. The studies were approved by the Internal Review Board of the College of Physicians and Surgeons of Columbia University.

##### CR/RANN Memory Tasks

all participants performed Selective Reminding Task (SRT) (Buschke & Fuld, 1974). Three memory measures were based on sub-scores of the SRT: the long-term storage sub-score, continuous long-term retrieval, and the number of words recalled on the last trial. The z-scores of each of the three measures were computed by subtracting the sample means followed by dividing by the sample standard deviation. The composite memory scores were computed as the average of the three z-scores.

#### HCPA

620 healthy participants with available cognitive data (age 36-100) from the lifespan Human Connectome Project Aging were included in this study (Bookheimer et al., 2019). The demographic information for the participants was presented in Table 1. HCPA excludes participants who have been diagnosed and treated for major psychiatric disorders (e.g., schizophrenia, bipolar disorder) or neurological disorders (e.g., stroke, brain tumors, Parkinson’s Disease). To be included in the current study, the following measurements had to be available: 1) T1-weighted MRI scans from 3T scanner, 2) years of education and recent occupation, and 3) Composite episodic memory score.

##### HCPA Memory Tasks

The cognitive and performance battery includes episodic memory measured by Picture Sequence Memory Test and Rey Auditory Verb al Learning Test (RAVLT). The z-scores of each of the three measures were computed by subtracting the sample means followed by dividing by the sample standard deviation. The composite memory scores were computed as the average of the three z-scores.

#### ADNI

941 subjects, including 417 normal control (CN), 378 mild cognitive impairment (MCI), and 146 Alzheimer’s disease (AD), were included from this study. The demographic information for the participants is presented in Table 1 and Table 3.

**Table 3.** Model performance for ADNI datasets by scanning manufacturers using random searching model

| Manufacturers |  | Tuning set (TL) | Test set before tuning (TL) | Test set after tuning (TL) | Tuning set (TLCO) | Test set (TLCO) |
| --- | --- | --- | --- | --- | --- | --- |
| **Siemens** | Rho | 0.6488 | 0.4589 | 0.5909 | 0.7836 | 0.5904 |
|  | MAE | 0.5349 | 0.8938 | 0.5163 | 0.5750 | 0.6244 |
| **GE** | Rho | 0.7813 | 0.5181 | 0.6558 | 0.5655 | 0.4636 |
|  | MAE | 0.4700 | 0.8461 | 0.5932 | 0.6816 | 0.6731 |
| **Philips** | Rho | 0.8264 | 0.3138 | 0.5785 | 0.5039 | 0.5238 |
|  | MAE | 0.3850 | 0.9285 | 0.7179 | 0.6406 | 0.6563 |

\*transfer learning (TL), and the hybrid (TLCO) approaches

Detailed inclusion and exclusion criteria for the ADNI study can be found at adni.loni.usc.edu. To be included in the current study, the following measurements had to be available: 1) T1-weighted MRI scans from 3T scanner, 2) years of education and recent occupation, and 3) Composite memory score (Crane et al., 2012). Written informed consent was obtained from all study participants according to the Declaration of Helsinki, and Ethical approval for data collection and sharing was given by the institutional review boards of the participating institutions in the ADNI.

##### ADNI Memory Tasks

ADNI memory was measured using modern psychometric approaches to analyze Rey Auditory Verbal Learning Test (RAVLT, 2 versions), AD Assessment Schedule – Cognition (ADAS-Cog, 3 versions), Mini-Mental State Examination (MMSE), and Logical Memory data. The composite scores were computed based on bifactor model (Crane et al., 2012). The computed data were downloaded from the ADNI website (UWNPSYCHSUM_03_26_20.csv).

## Image Procedures

### Neuroimaging data acquisition

#### RANN

Structural MRI scans were acquired on a 3.0T Philips Achieva scanner. T1-weighted MPRAGE scan was acquired with a TE/TR of 3/6.5 ms and Flip Angle of 8°, inplane resolution of 256 × 256, field of view of 25.4 × 25.4 cm, and 165–180 slices in axial direction with slice-thickness/gap of 1/0 mm.

#### HCPA

Structural MRI scans were acquired from all sites using 3T Siemens Prisma scanner, and 32-channel Prisma head coil. T1-weighted images were acquired with 3D multiecho magnetization prepared rapid gradient echo (MEMPRAGE) at 0.8 mm isotropic resolution (Harms et al., 2018). Other parameters include: TR/TI = 2500/1000, TE = 1.8/3.6/5.4/7.2 ms, flip angle of 8 deg, FOV of 256 × 240 × 166 mm with a matrix size of 320 × 300 × 208 slices, water excitation employed for fat suppression (to reduce signal from bone marrow and scalp fat), and up to 30 TRs allowed for motion-induced reacquisition.

#### ADNI

Structural MRI scans were acquired from all sites using 3T Philips, GE, and Siemens scanners. Since the acquisition protocols were different for each scanner, an image normalization step was performed by the ADNI. The imagining sequence was a 3-dimensional sagittal part magnetization prepared of rapid gradient-echo (MPRAGE). This sequence was repeated consecutively to increase the likelihood of obtaining at least one decent quality of MPRAGE image. Image corrections involved calibration, geometry distortion, and reduction of the intensity of non-uniformity applied on each image by the ADNI. More details concerning the sMRI images is available on the ADNI homepage (http://adni.loni.usc.edu/methods/mri-tool/mri-analysis/).

### Neuroimaging data processing

Each subject’s structural T1 scan was reconstructed using FreeSurfer v7.1.1 (http://surfer.nmr.mgh.harvard.edu/). The accuracy of FreeSurfer’s subcortical segmentation and cortical parcellation (Fischl et al., 2002) has been reported to be comparable to manual labeling. All T1 images went through an automated quality control through MRIQC (Esteban et al., 2017). For the multiple available T1 images at the same visit, we selected the images with the best quality for further analysis. For all images that passed quality check, crosssectional image processing was performed using FreeSurfer Version 7.1.1 (https://surfer.nmr.mgh.harvard.edu/). Region of interest (ROI)-specific cortical thickness and volume measures were extracted from the automated anatomical parcellation using the Desikan-Killiany Atlas (Desikan et al., 2006) for cortical and aseg atlas for subcortical ROIs. To test the robustness of all the models (supplementary material), we also used an alternative Destrieux atlas (Destrieux, Fischl, Dale, & Halgren, 2010).

### Brain memory prediction model

The memory prediction model was trained using the RANN dataset. An overview of the transfer learning method is presented in Figure 1. First, the RANN dataset was split into the training set (70%) and test set (30%) using a conditionally random method. The distributions of age and sex in the two sets were statistically identical. Cascade neural network models with all regional cortical thickness and volume from FreeSurfer as inputs were used to train the RANN dataset for memory prediction. The cascade neural network is a feedforward neural network involving connections from the input and every previous layer to the subsequent layer (Figure 2). The advantage of the model is that it accommodates the nonlinear relationship between input and output. It has been shown to outperform the other common classical machine learning approaches for brain residual-based analysis, and more flexible and efficient to implement the transfer learning framework compared to other approaches. (Chen et al., 2020). The hyperparameters of the model, including numbers of hidden layers, numbers of neurons, penalty of regularization and types of activation function, were optimized through random search. The loss function of model optimization was specified as mean square error function optimized using gradient descent algorithm with an adaptive learning rate and constant momentum. A 10-fold cross-validation procedure was conducted within the training set to estimate the memory prediction model performance. To quantify model performance, metrics including Pearson’s correlation coefficient (rho), mean absolute error (MAE) and Cohen’s *f^2^* between the predicted and true memory were calculated.

**Figure 1.**
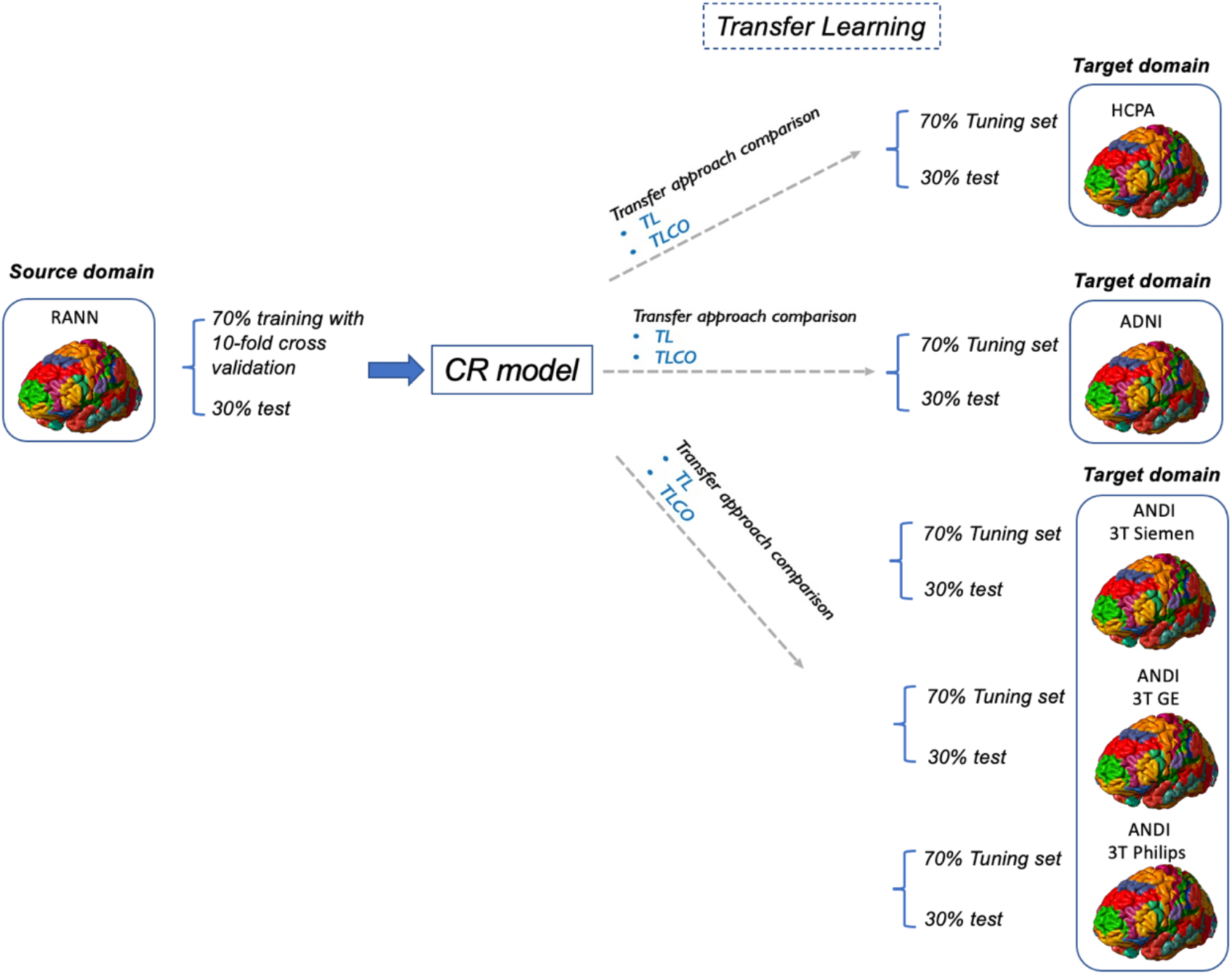
Overview of transfer learning methods

**Figure 2:**
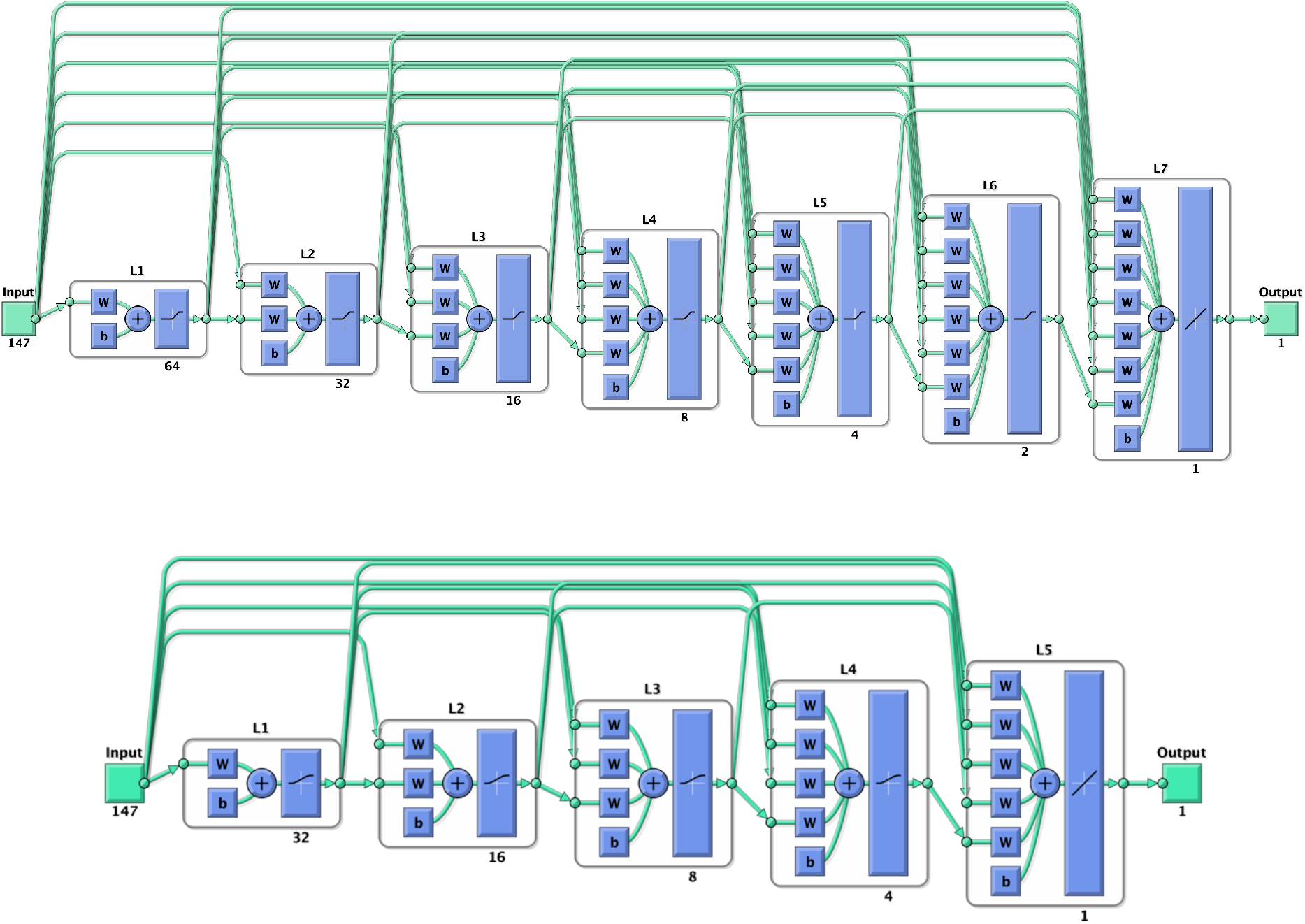
the initial cascade neural network model used to train CR model (top). This network trained using RANN dataset has seven layers (L1 to L7). The number under each layer represents the number of neurons in that layer. The first layer has a weight coming from the input and each subsequent layer has weight coming from the inputs with all previous layers. The last layer is the network output, called as output layer. The output layer is also connected directly with the input layer beside with hidden layer. The optimized CNN model after random search (bottom) included 5 layers.

### Transfer learning

To transfer the T1 imaging-based model from source domain (using RANN) to the target domain (HCPA or ADNI), we first randomly divided the whole HCPA or ADNI dataset into the tuning pool and test sets (tuning set 70%, test set 30%). The subset of the tuning pool was randomly selected to re-train the pre-trained model.

In the refined optimization procedure of the transfer learning, the optimal tuning sample, the regularization ratio (0 to 1), the loss function (i.e. mean square error), and the choice which layers were frozen (if the layer of the pre-trained model was frozen, the parameters in that layer were not updated in the fine-tuning process) were tested. The transfer learning process was optimized using an agile optimization process because it facilitated rapid prototyping and broad searching. After the tuning procedure, the transferred model was applied to the test set for model performance evaluation. We compared the performance of transfer learning approach with the TLCO, which is used for re-training the pre-trained memory prediction model by using a combination of the tuning and training sets with the site indicator. We optimized the tunning process by adopting an agile optimization method that exploit a time-saving optimizer called scaled conjugate gradient (SCG) algorithm for fast optimization and the hyperparameter settings emulated as those of the training process in the target domain. We compared the performance of the optimized transfer learning with tuning procedure with the model applied pre-trained model without tuning. Since ADNI data was collected from multiple sites and multiple scanners, for the secondary analysis, we applied the transfer learning by the scanner manufacturers: GE, Siemens and Philips. The data in the three target domains were divided in to the tuning-pool and test sets (Siemens: tuning pool N=377, test set N=160; GE: tuning pool N=164, test set N=68; Philips: tuning pool N=124, test set N=48). Then, transfer learning was performed in the same pattern separately for three datasets. The transfer learning with cotrain (TLCO) method was used to compare the performance with transfer learning. TLCO integrated both training set from source domain and tunning set from target domain to tune the pre-trained CNN model. The TLCO approach accounts for intersite differences through statistical variance analysis. It employs statistical models to regress out site-specific differences by using statistical covariates. This approach requires the source domain data to be accessible and the data size from different sites to be balanced. The code of the transfer learning is available at https://github.com/XiZhu-CU/Transfer-Learning-Submission.

### Quantification of cognitive reserve

After establishing the memory prediction model, a person’s predicted memory performance could be obtained. Structural brain features along with age and sex were included in the model as predictors (Reed et al., 2010). Race was not included in the model as a predictor because more than 93% of our targeted sample (ADNI) is non-Hispanic white. The impact of race on the model performance is presented in supplementary material. In addition, the estimated intracranial volume (eTIV) was extracted from each subject and used as a predictor. Cognitive reserve was quantified by residuals which subtract the predicted memory from the true memory. To validate our brain-based CR quantification, we performed correlation analyses between the residuals and several proxies of CR including education, occupation and IQ. For RANN and ADNI, we used National Adult Reading Test (NART) IQ, which reflects the crystallized intelligence. Occupational attainment variables (data, people, things) reflect the specific demands of an occupation. All RANN and ADNI findings were corrected for multiple comparison at p<0.01 (5 measures). Similarly, for HCPA, the NIH Toolbox was administrated provided the Crystallized composite scores which reflects the intelligence. The Crystallized Composite score is derived from performance on the Reading Recognition and the Picture Vocabulary tasks (Heaton et al., 2014). The HCPA findings were corrected for multiple comparison at p<0.025 (2 measures).

## Results

### Demographic Characteristics

Demographic and clinical characteristics are presented in Table 1. All three datasets significantly differed in age, sex, and education. Participants were older in ADNI compared with RANN and HCPA. Education was higher in HCPA subjects, compared with RANN or ADNI. IQ was not significantly different between CRNN and ADNI.

### Training memory prediction modeling in the RANN dataset

The cascade neural network model (Figure 2) using 10-fold cross-validation on the RANN training set demonstrated significant linear correlation between true and predicted memory based on the chosen T1 cortical thickness and volume predictors for both training set (rho=0.6076, MAE=0.5856, Cohen’s ***f^2^***=0.58) and independent test set (rho=0.3886, MAE=0.6980, Cohen’s ***f^2^***=0.18) (Figure 3). After random search, the model performance improved in training set (rho=0.5578, MAE=0.5792, Cohen’s ***f^2^***=0.45) and test set (rho=0.3963, MAE=0.6888, Cohen’s ***f^2^***=0.19).

**Figure 3.**
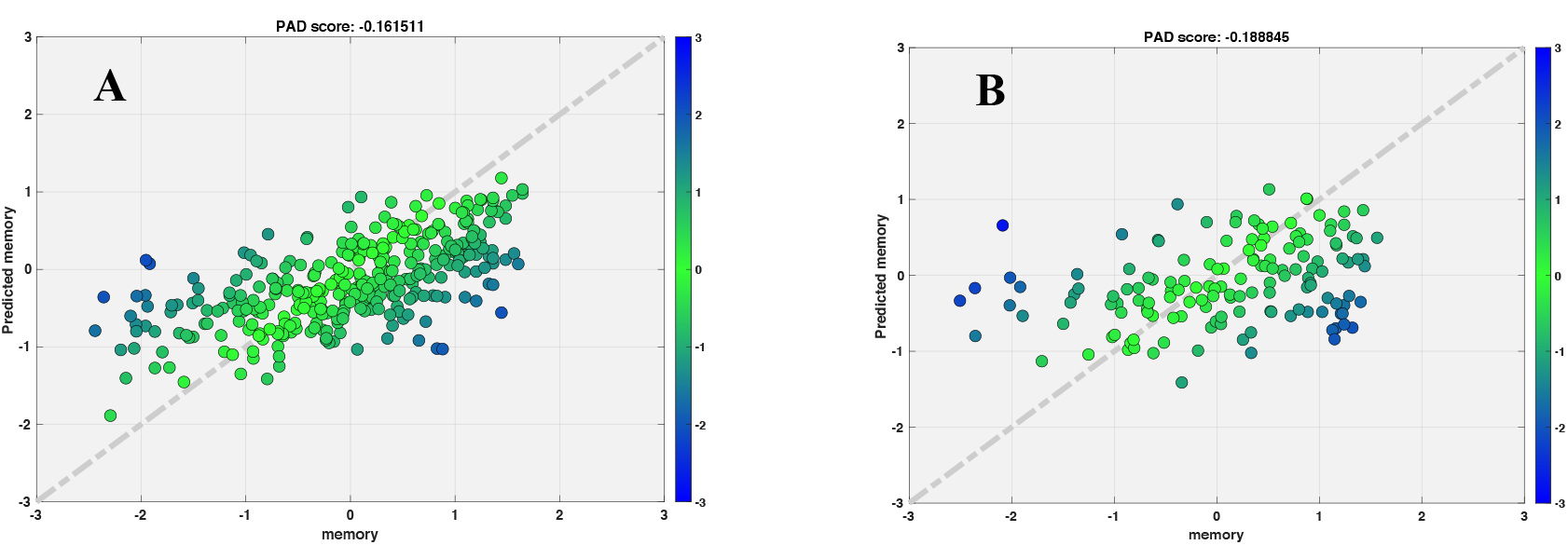
Scatter plot for true memory (x axis) against predicted memory (y axis) in RANN dataset after random search. A) Training set; B) Test set

There was significant correlation of NART IQ with residuals for training set (NART IQ: rho=0.154, p-value =.004, Cohen’s ***f^2^***=0.01). There was significant correlation between the residuals and both NART IQ (NART IQ: rho=0.169, p-value =.003, Cohen’s ***f^2^***=0.03) and education (rho=0.2069, p-value=0.01, Cohen’s ***f^2^***=0.04) for test set. Residuals were not associated with data, people or things.

### Transfer learning to HCPA

The best model trained using RANN dataset (pre-trained model) was used in this analysis. First, we tuned the model using tuning set from target domain (HCPA). We found linear correlation and low MAE between true and predicted memory for tuning set (rho=0.4909, MAE=0.4101, Cohen’s ***f^2^***=0.32) and test set (rho=0.4062, MAE=0.4107, Cohen’s ***f^2^***=0.20). When we directly applied pre-trained model without tuning, the performance dropped in test set (rho=0.3099, MAE=0.5358, Cohen’s ***f^2^***=0.11) (Figure 4). Second, the transfer learning with cotrain (TLCO) approach uses both training set from source domain (RANN) and tunning set from target domain (HCPA) to further tune the pretrained model. The TLCO performed comparable with the transfer learning approach (Tuning set: rho=0.3872, MAE=0.4318, Cohen’s ***f^2^***=0.18; Test set: rho=0.4474, MAE=0.3867, Cohen’s ***f^2^***=0.25).

**Figure 4.**
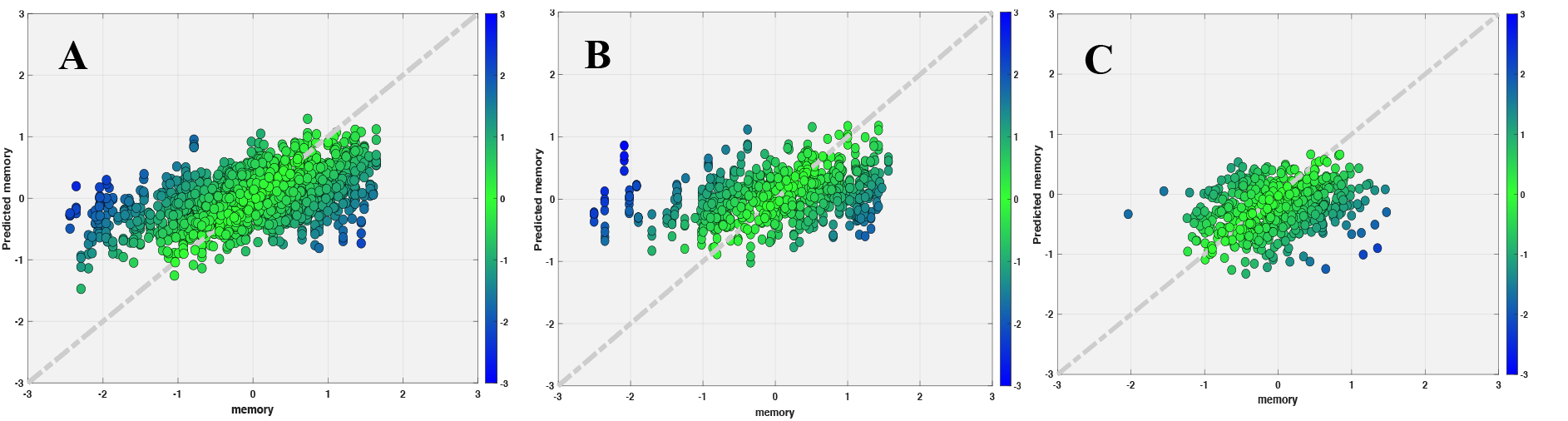
Scatter plot for true memory (x axis) against predicted memory (y axis) in HCPA dataset, while applied pretrained model from RANN. A) Tuning set using HCPA data; B) Test set after tuning using HCPA data; C) Test set if applying the pretrained model directly

There was significant correlation of both IQ and education with residuals of the transfer learning model for both tuning set (IQ: rho=0.227, p-value <.001, Cohen’s ***f^2^***=0.05; education: rho=0.255, p-value=0.0015, Cohen’s ***f^2^***=0.07); IQ: rho=0.3612, p-value <.001, Cohen’s ***f^2^***=0.15; education: rho=0.2798, p-value<.001, Cohen’s ***f^2^***=0.09).

### Transfer learning to ADNI

#### 1. Primary analysis

We found strong linear correlation and low MAE between true and predicted memory for tuning set (rho=0.7385, MAE=0.4935, Cohen’s ***f^2^***=1.2) and test set (rho=0.7117, MAE=0.5435, Cohen’s ***f^2^***=1.03). When we directly applied the pre-trained model without tuning, performance dropped in test set (rho=0.5485, MAE=0.9259, Cohen’s ***f^2^***=0.43) (Figure 5). The TLCO performed comparable with the transfer learning approach (Tuning set: rho=0.7187, MAE=0.5158, Cohen’s ***f^2^***=1.1; Test set: rho=0.6684, MAE=0.5967, Cohen’s ***f^2^***=0.81).

**Figure 5.**
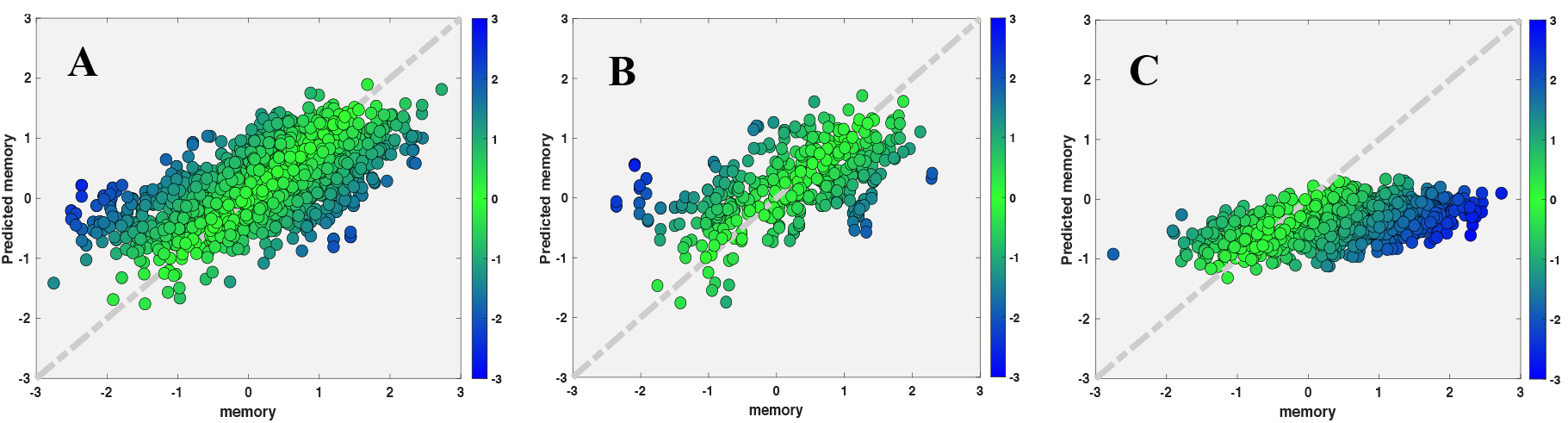
Scatter plot for true memory (x axis) against predicted memory (y axis) in ADNI dataset, while applied pretrained model from RANN. A) Tuning set; B) Test set after tuning; C) Test set if applying the pretrained model directly

There was significant correlation between IQ, education, and residuals of the transfer learning model for both tuning set (IQ: rho=0.2025, p-value < .001, Cohen’s ***f^2^***=0.04; education: rho=0.1698, p-value=0.0032, Cohen’s ***f^2^***=0.03) and test set (IQ: rho=0.366, p-value < .001, Cohen’s ***f^2^***=0.15; education: rho=0.255, p-value < .001, Cohen’s ***f^2^***=0.07).

We further assessed the correlation of IQ and education with residuals separately within each diagnosis group (CN, MCI and AD). Correlations of NART IQ and education with residuals are presented in Table 4. Significant correlation between IQ and residuals was found in all three groups, while the significant correlation between education and residuals was found in CN and MCI.

**Table 4.**
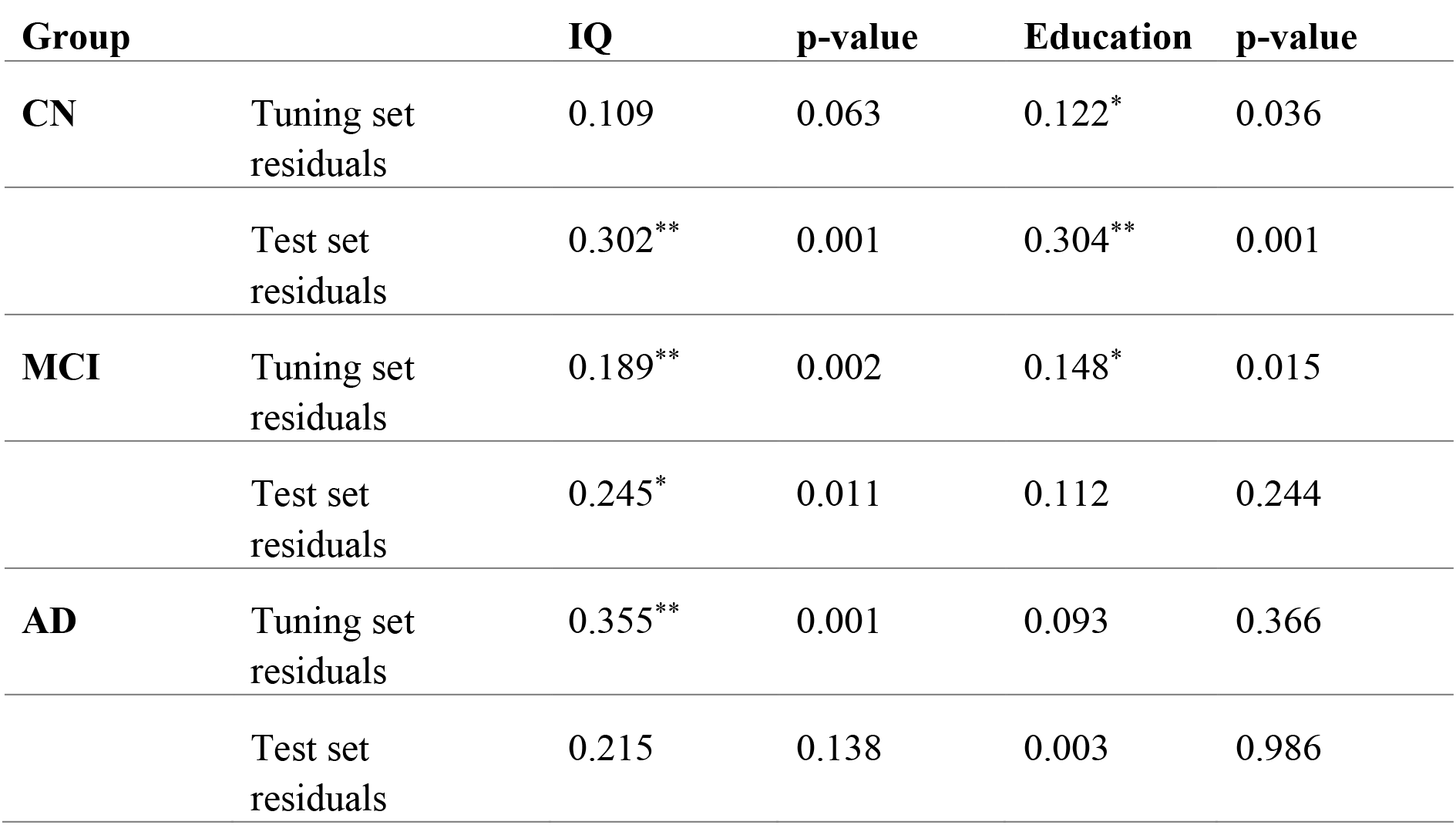
Pearson’s correlation coefficient between IQ, education and residuals by group diagnosis in ADNI dataset

#### 2. Secondary analysis

We found strong linear correlation and low MAE between true and predicted memory in the Siemens, GE and Philips respectively for tuning set (Siemens: rho=0.6488, MAE=0.0.5349; GE: rho=0.7813, MAE=0.4700, Cohen’s *f^2^*=1.57; Philips: rho=0.8264, MAE=0.3850, Cohen’s ***f^2^***=2.15) and test set (Siemens: rho=0.5909, MAE=0.5163, Cohen’s ***f^2^***=0.54; GE: rho=0.6558, MAE=0.5932, Cohen’s ***f^2^***=0.75; Philips: rho=0.5785, MAE=0.7179, Cohen’s ***f^2^***=0.50). Using the transfer learning approach, the performance of the RANN pre-trained model could be reproduced in each target domain with a smaller amount of tuning data (Figure 6). The transfer learning approach always outperformed the TLCO. The results are shown in Table 3.

**Figure 6.**
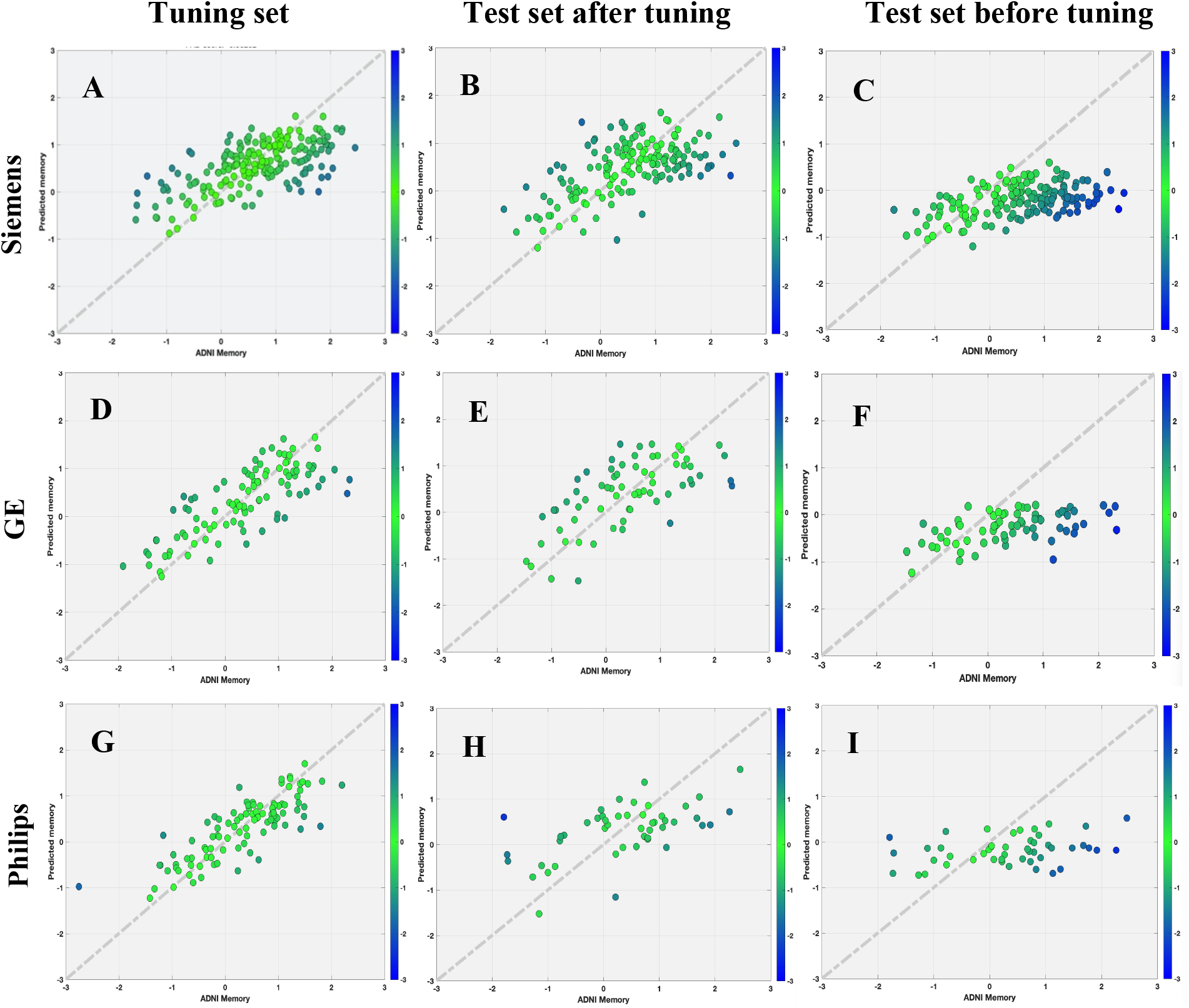
Scatter plot for true memory (x axis) against predicted memory (y axis) in ADNI dataset by scanner types using random searching models.

Significant and positive correlations between NART IQ, education and residuals were demonstrated in both tuning and test sets (Table 5).

**Table 5.** Pearson’s correlation coefficient between IQ, education and residuals by scanner types in ADNI dataset

| **Manufacturer** |  | **IQ** | **p-value** | **Education** | **p-value** |
| --- | --- | --- | --- | --- | --- |
| **Siemens** | Tuning set residuals | 0.2364** | < .001 | 0.2171** | 0.0020 |
|  | Test set residuals | 0.1933* | 0.0143 | 0.1858* | 0.0186 |
| **GE** | Tuning set residuals | 0.2700** | 0.0085 | 0.1999* | 0.0461 |
|  | Test set residuals | 0.0886 | 0.4791 | 0.3130* | 0.0094 |
| **Philips** | Tuning set residuals | 0.1996* | 0.0488 | 0.2808** | 0.0047 |
|  | Test set residuals | 0.5001** | <.001 | 0.3424** | 0.0172 |

## Discussion

In this study, we built a deep learning model to quantify the CR as residual variance in memory performance using the sMRI data from a healthy lifespan cohort (age 20-80). Importantly, our study demonstrates that the pre-trained model constructed using the healthy lifespan data (RANN) from a single-site and a single sequence was able to generalize to two target datasets acquired with different age ranges, imaging protocols, and clinical status. These included healthy lifespan Human Connectome Project-Aging cohort (HCPA) and older MCI and demented participants from Alzheimer’s Disease Neuroimaging Initiative (ADNI) across different scanner types. By tuning the models with relatively small sample sizes and the same T1 brain features, optimal transferred models were obtained with satisfactory prediction performance in both target cohorts. The estimated CR was also validated by showing significant correlation with CR proxies such as education and IQ across all three datasets.

We found that the cascade neural network (CNN) model trained on the RANN data demonstrated a linear correlation between true and predicted memory based on the T1 cortical thickness and volume predictors. The sMRI-based measure of CR was associated with CR the proxy measures of education and IQ. Previous studies have used sMRI from older healthy subjects (Sole-Padulles et al., 2009), older MCI, or patients with AD to quantify CR (van Loenhoud et al., 2017). However, patients with neurological diseases with aberrant cognition may lead to bias for the quantification of CR. We first demonstrated that using lifespan data of healthy individuals, enabled good quantification of cognitive performance. It is worth noting that the performance achieved by our model is also comparable to that of previous studies applying residual approaches on quantifying CR (Vieira, Pinaya, & Mechelli, 2017).

Second, to test the generalizability of the sMRI-based deep learning model, this study utilized the transfer learning approach to fine-tune the pre-trained deep learning model to an independent, healthy lifespan HCPA data. The transfer learning approach is an efficient and stabilized way to generalize the T1 imaging-based memory prediction model. Compared with the TLCO, the tuning methods with the transfer learning approach always provided lower MAE and a stronger correlation between the actual and predicted memory in all results.

Third, the model not only could generalize from healthy lifespan data to an independent healthy lifespan HCPA dataset, but also to an older demented participants from ADNI using transfer learning. Although the three datasets administrated different tests to assess memory, by tunning the models with relatively small sample size, prediction performance of the models were relatively comparable. Moreover, the models were robust across different scanners. When conducting retrospective multi-center imaging studies, such as ADNI, or applying models trained on one site to another, heterogeneous MRI data from different scanner hardware, and acquisition protocols will pose challenges in the evaluation and generalization of these trained models. Structured programs aimed at standardizing and harmonizing MRI acquisition in research settings (Weiner et al., 2017). However, data obtained in these selected frameworks might not be representative of real-world populations. In our work, CNN was trained, tested using the RANN dataset, then using transfer learning to fine-tune and test in another two datasets obtained by different MR protocols and scanners to capture the full spectrum of heterogeneity among data and provide a less dataset-specific approach. Through further training iterations, the pre-trained CNN network adjusted for data bias stemming from the differences in acquisition and reconstruction between different scanners. In fact, our approach overcomes the caveats of previous work, which obtained data from single-center datasets leading to a limited reproducibility of findings (Wen et al., 2020).

In the current study, we used a standard pipeline to process the raw MRI image and extracted the cortical thickness and volume measures from T1-weighed MRIs. Our ROI-based approach shown promising and robust results for the given sample sizes. Further deep learning studies with larger sample size may also consider using voxel-wise whole brain based approach as input. Moreover, despite progress on the interpretability of deep learning, deep neural networks are still considered, to a large extent, as black boxes, due to the difficulty of interpreting their inner networks. For example, even when an model allows detection of patients from controls with high levels of accuracy, it can be difficult to establish the specific features that informed the classification decision (Cruz-Roa, Arevalo Ovalle, Madabhushi, & Gonzalez Osorio, 2013). However, our focus of this study was to better predict the memory measures, further studies may develop more interpretable deep learning models to better understand the underlying neural mechanism. Lastly, we only used sMRI to assess the feasibility for CR estimation across three studies. Future studies should consider adding other MRI modalities, such as, diffusion tensor imaging (DTI), PET, and CSF biomarkers together with sMRI to improve the power of prediction as well as the accuracy of the residual in estimating CR.

## Conclusions

In conclusion, we have shown the general feasibility of using deep learning to quantify cognitive reserve by leveraging lifespan healthy data. Our findings showed that brain/cognitive function across lifespan provided good brain-based quantification of CR. Moreover, transfer learning shows promises for building robust models that can be fine-tuned and generalized to independent healthy lifespan cohort and in patients with Alzheimer’s disease, also robust across different scanners with different acquisition parameters. The residuals (CR) were significantly associated with NART IQ and education across different cohorts. The transfer learning method is applicable to various brain diseases or CR proxies and may flexibly incorporate different imaging modalities making it a promising tool for scientific and clinical purposes.

## Data Availability

All data examined in the manuscript are available upon request in deidentified format.

