## supplemental material for "Transfer Learning for Cognitive Reserve Quantification"

### Supplementary information

#### 1. Test the impact of race on model performance

**Training memory prediction modeling in the RANN dataset including race as a predictor**

The cascade neural network model (Figure S1) using 10-fold cross-validation on the RANN training set demonstrated significant linear correlation between true and predicted memory based on the chosen T1 cortical thickness and volume predictors for both training set and independent test set. After random search, the model performance in training set (rho=0.5193, MAE=0.633) and test set (rho=0.4019, MAE=0.6910).

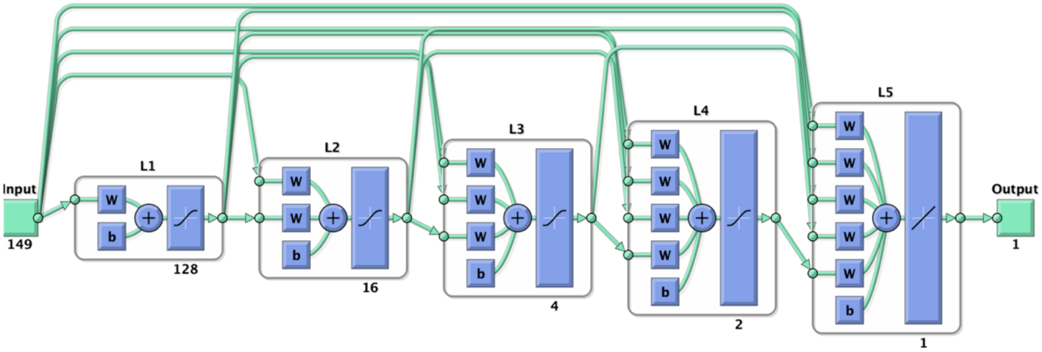


Figure S1: The optimized CNN model after random search included 5 layers.

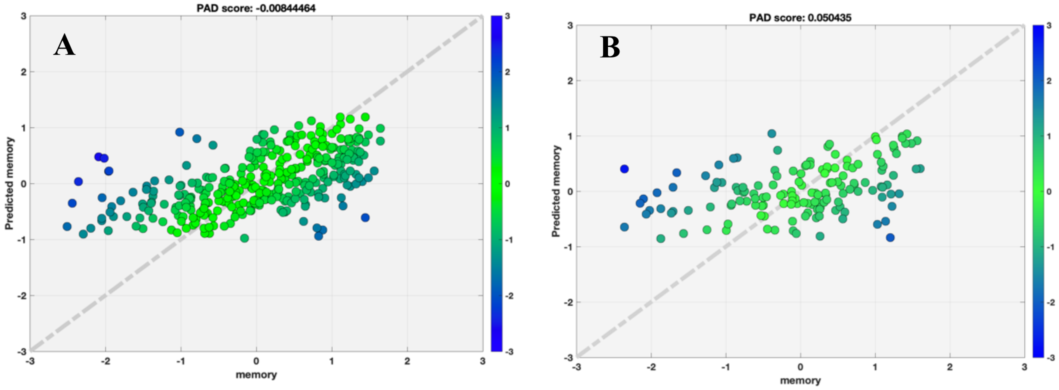


Figure S2: Scatter plot for true memory (x axis) against predicted memory (y axis) with race using Desikan-Killiany Atlas in RANN dataset after random search.
A) Training set; B) Test set

**Transfer learning to HCPA**

We found linear correlation and low MAE between true and predicted memory for tuning set (rho=0.1921, MAE=0.4509) and test set (rho=0.4059, MAE=0.4067) (Figure S4). When we directly applied pre-trained model without tuning, the performance remains similar in test set (rho=0.4087, MAE=0.4024).

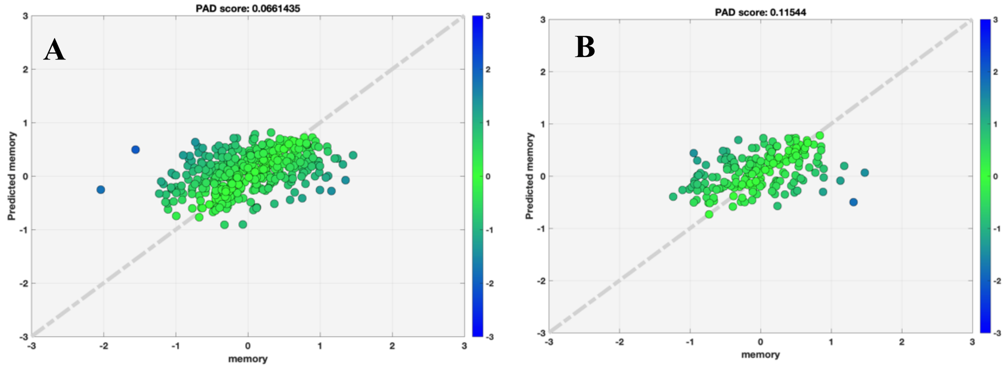


Figure S4: Scatter plot for true memory (x axis) against predicted memory (y axis) with race using Desikan-Killiany Atlas in HCPA dataset after random search.
A) Training set; B) Test set

**Transfer learning to ADNI**

We found strong linear correlation and low MAE between true and predicted memory for tuning set (rho=0.7364, MAE=0.5150) and test set (rho=0.6406, MAE=0.5650) (Figure S5). When we directly applied the pre-trained model without tuning, performance dropped in test set (rho=0.3902, MAE=0.9316).

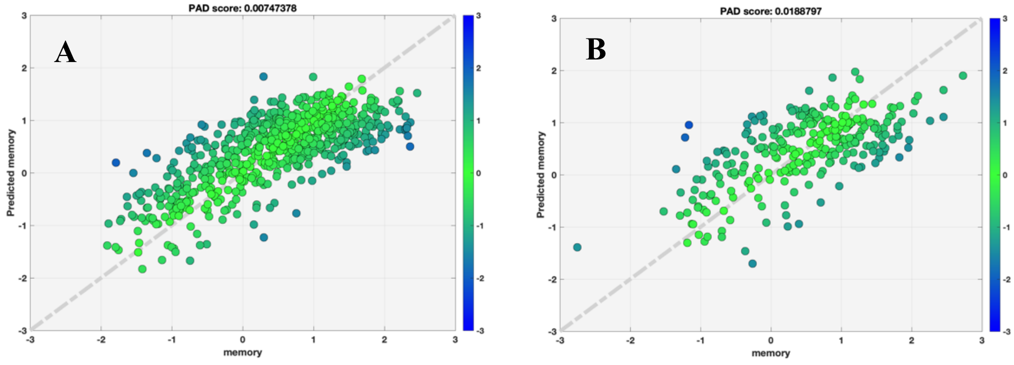


Figure S5: Scatter plot for true memory (x axis) against predicted memory (y axis) with race using Desikan-Killiany Atlas in ADNI dataset after random search.
A) Training set; B) Test set

2. **Test the impact of atlas on model performance (using Destrieux atlas in the following analysis)**

**Training memory prediction modeling in the RANN dataset including race as a predictor**

The cascade neural network model (Figure S6) using 10-fold cross-validation on the RANN training set demonstrated significant linear correlation between true and predicted memory based on the chosen T1 cortical thickness and volume predictors for both training set and independent test set. After random search, the model performance using Destrieux in training set is (rho=0.5352, MAE=0.6493) and test set (rho=0.4048, MAE=0.6941) (Figure S7).

There was significant correlation of NART IQ with residuals for training set (NART IQ: rho=0.168 with p-value =.0018). There was no significant correlation between the residuals and education for test set. Residuals were not associated with data, people or things.

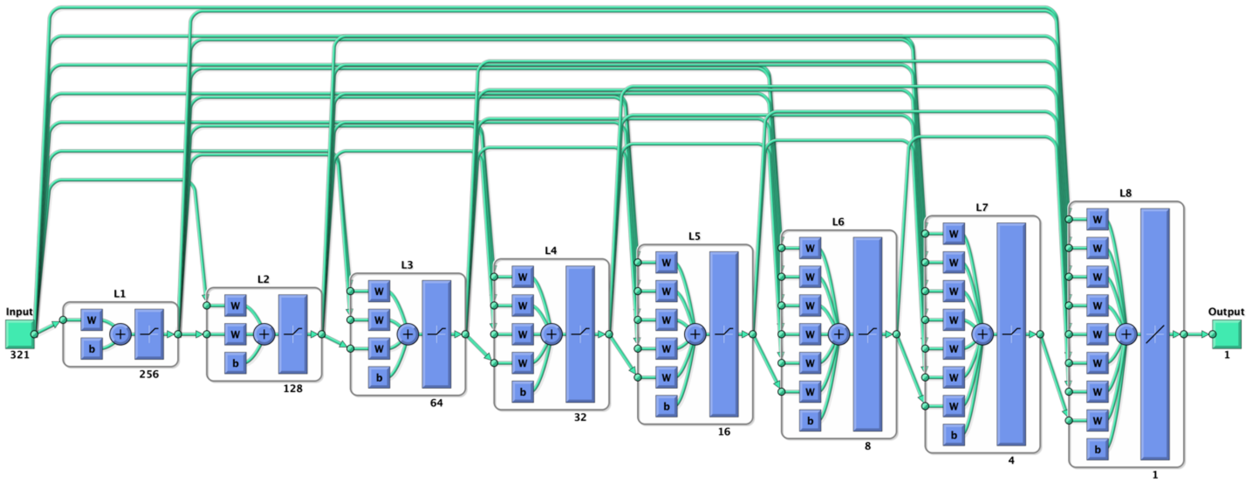


Figure S6: The optimized CNN model using Destrieux atlas after random search included 8 layers.

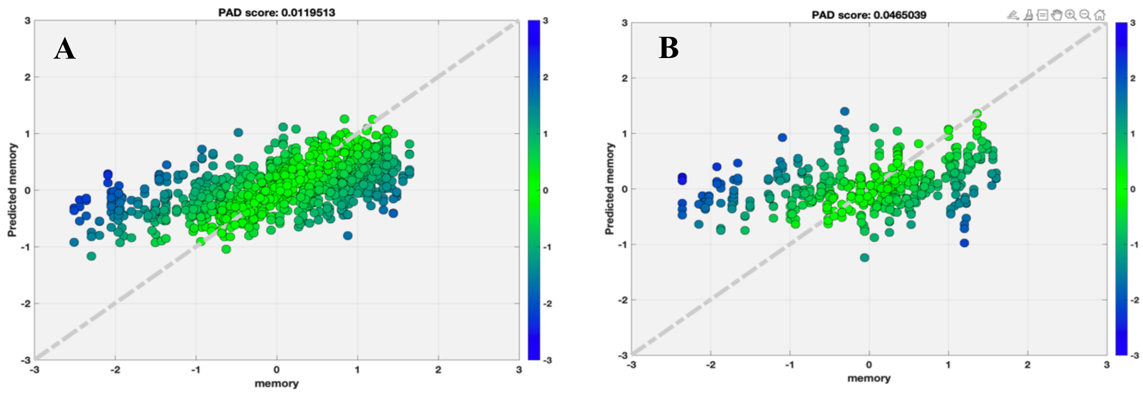


Figure S7: Scatter plot for true memory (x axis) against predicted memory (y axis) with race using Destrieux Atlas in RANN dataset after random search.
A) Training set; B) Test set

**Transfer learning to HCPA using Destrieux Atlas**

The best model trained using RANN dataset (pre-trained model) was used in this analysis. We found linear correlation and low MAE between true and predicted memory for tuning set (rho=0.2448, MAE=0.4545) and test set (rho=0.4286, MAE=0.3806). When we directly applied pre-trained model without tuning, the performance dropped in test set (rho=0.3053, MAE=0.5187) (Figure S8).

There was significant correlation of both IQ and education with residuals for both tuning set (IQ (fluid): rho=0.255 with p-value <.001; IQ (Crystallized): rho=0.159 with p-value <.001; education: rho=0.12 with p-value=0.01) and test set (IQ (Crystallized): rho=0.256 with p-value <.001; education: rho=0.205 with p-value<.001).

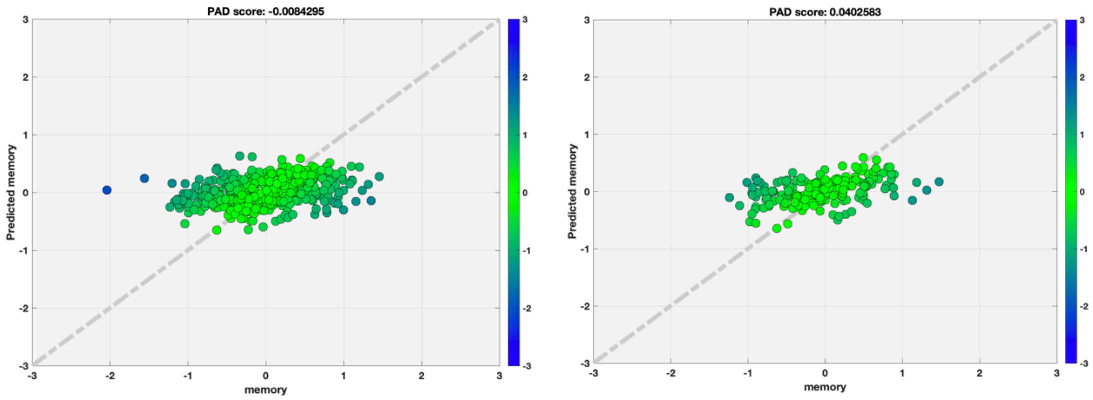


Figure S8: Scatter plot for true memory (x axis) against predicted memory (y axis) with race using Destrieux Atlas in HCPA dataset after random search.

A) Training set; B) Test set

##### Transfer learning to ADNI using Destrieux Atlas

We found strong linear correlation and low MAE between true and predicted memory for tuning set (rho=0.6912, MAE=0.5592) and test set (rho=0.6200, MAE=0.5684). When we directly applied the pre-trained model without tuning, performance dropped in test set (rho=0.2375, MAE=0.8570) (Figure S9).

There was significant correlation between IQ, education, and residuals for both tuning set (IQ: rho=0.3062 with p-value < .001; education: rho=0.1999 with p-value<.001) and test set (IQ: rho=0.22 with p-value < .001; education: rho=0.20 with p-value < .001).

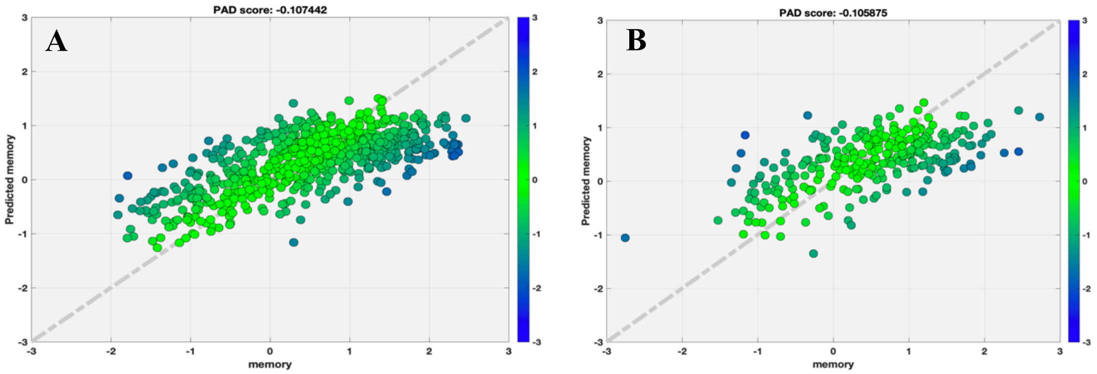


Figure S9: Scatter plot for true memory (x axis) against predicted memory (y axis) with race using Destrieux Atlas in ADNI dataset after random search.
A) Training set; B) Test set

##### 3. Test the impact of tuning and testing split on target domain on model performance (using Desikan-Killiany atlas)

We tested the impact of the size of tuning set on target model performance on HCP dataset (Figure S10). There was only slightly variability on the model performance (r and MAE) using 10%~50% test set. Once we used less than 50% data for tuning and more than 50% data for testing, then the model performance dropped in HCPA dataset.

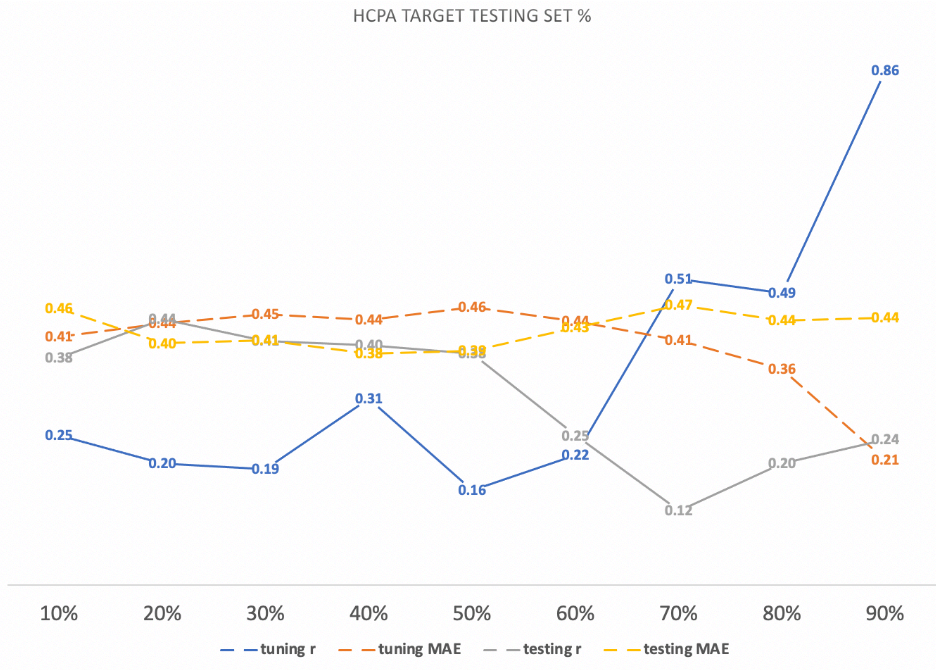


Figure S10: target domain (HCPA) tuning testing split vary from 10% testing (left) to 90% testing (right)

We tested the impact of the size of tuning set on target model performance on ADNI dataset (Figure S10). There was only slightly variability on the model performance (r and MAE) using 10%~80% test set. Once we only used 10% data for tuning and 90% data for testing, then the model performance dropped in ADNI dataset.

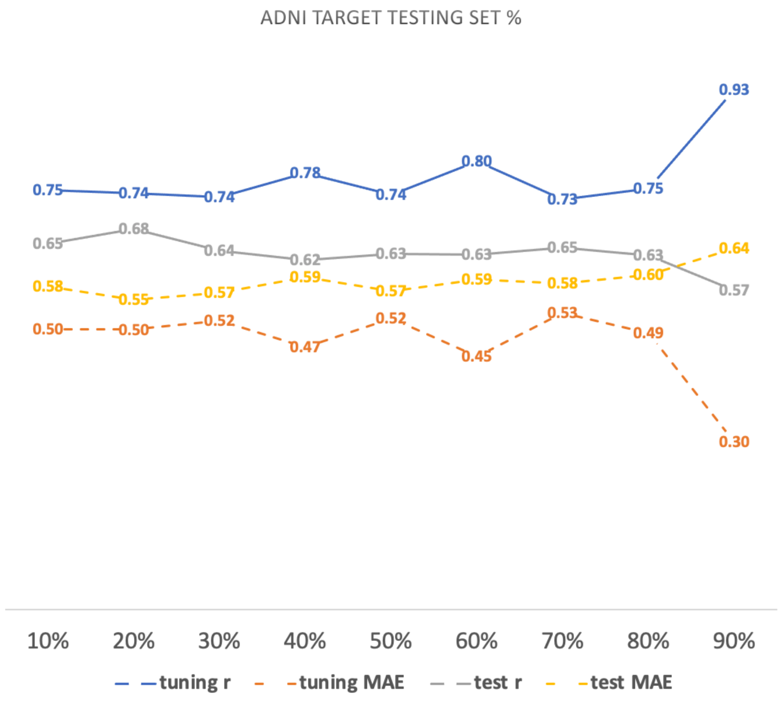


Figure S11: target domain (ADNI) tuning testing split vary from 10% testing (left) to 90% testing (right)

Since the ADNI dataset is a more heterogenous dataset including CN, MCI and AD, and covers a wide range of the memory measure across all subjects (Figure S12 A). Therefore, this target domain requires less tuning dataset (as low as 20% tuning set) for fine tune the transfer learning model. On the other hand, the HCPA dataset is a more homogenous dataset including only CN, and covers relatively narrower range of the memory measure across all subjects (Figure S12 A). This target domain requires at least 50% tuning dataset for fine tune the transfer learning model.

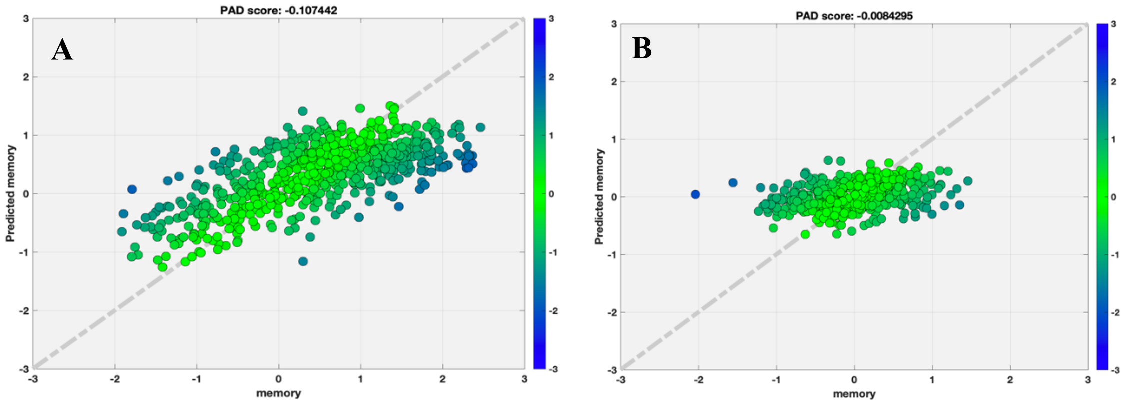

Figure S12: Comparison of the HCPA and ADNI datasets.
